# Sapphire-Based Optrode for Low Noise Neural Recording and Optogenetic Manipulation

**DOI:** 10.1101/2024.08.15.608194

**Authors:** Yanyan Xu, Ben-Zheng Li, Xinlong Huang, Yuebo Liu, Zhiwen Liang, Xien Yang, Lizhang Lin, Liyang Wang, Yu Xia, Matthew Ridenour, Yujing Huang, Zhen Yuan, Achim Klug, Sio Hang Pun, Tim C. Lei, Baijun Zhang

**Author notes:** indicates equal contribution (Yanyan Xu, Ben-Zheng Li). **All correspondence and materials requests should be addressed to:** Baijun Zhang, PhD, State Key Laboratory of Optoelectronic Materials and Technologies, School of Electronics and Information Technology, Sun Yat-sen University, Guangzhou, 510275, China.

## Abstract

Electrophysiological recordings of neurons in deep brain regions using optogenetic stimulation are essential to understanding and regulating the role of complex neural activity in biological behavior and cognitive function. Optogenetic techniques have significantly advanced neuroscience research by enabling the optical manipulation of neural activities. Because of the significance of the technique, constant advancements in implantable optrodes that integrate optical stimulation with low-noise, large-scale electrophysiological recording are in demand to improve the spatiotemporal resolution for various experimental designs and future clinical applications. However, robust and easy-to-use neural optrodes that integrate neural recording arrays with high-intensity light emitting diodes (LEDs) are still lacking. Here, we propose a neural optrode based on Gallium Nitride (GaN) on sapphire technology, which integrates a high-intensity blue LED with a 5×2 recording array monolithically for simultaneous neural recording and optogenetic manipulation. To reduce the noise interference between the recording electrodes and the LED, which is in close physical proximity, three metal grounding interlayers were incorporated within the optrode, and their ability to reduce LED-induced artifacts during neural recording was confirmed through both electromagnetic simulations and experimental demonstrations. The capability of the sapphire optrode to record action potentials has been demonstrated by recording the firing of mitral/tuft cells in the olfactory bulbs of mice in vivo. Additionally, the elevation of action potential firing due to optogenetic stimulation observed using the sapphire probe in medial superior olive (MSO) neurons of the gerbil auditory brainstem confirms the capability of this sapphire optrode to precisely access neural activities in deep brain regions under complex experimental designs.

## INTRODUCTION

Recording action potentials fired by neurons is a well-used technique in neuroscience research to study the functions and working principles of neuronal circuits in the brain. Optogenetics, which utilizes viral vectors to express opsins in neurons, has become an important experimental tool for stimulating or inhibiting neural activities by controlling ionic flows across neuronal membranes using light^1–4^. To allow simultaneous recording of action potentials and optogenetic manipulation of neuronal activities, investigators glue optical fibers onto neural recording probes to create “optrodes” which can simultaneously record neural activity and deliver light to target brain nuclei. However, custom-making optrodes for experimental use is associated with several challenges which can lead to failed experiments and unnecessary animal use. Aligning the fiber tip to the recording site is difficult, and light emitted from the fiber tip often misses target neurons, resulting in a lack of opsin stimulation and subsequent experimental failures. Additionally, the addition of a fiber to the recording electrode enlarges the insertion diameter of the optrode, which can cause more brain tissue damage during probe insertion^5–7^. There have been previous engineering attempts to overcome the issues with optrode design and manufacturing. Waveguide-coupled silicon electrodes have been manufactured to couple light emitted from an external laser or light emitting diode (LED) to the target nucleus^8–11^. However, these waveguide-coupled optrodes rely on light generated from external sources, such as lasers or LEDs. Not only can the coupling efficiency between the fiber and the light source be low—as low as a few percent—but the tethering fiber can also be fragile, significantly limiting the movements of the animals, which is problematic when behavioral responses are important^12,13^. Therefore, developing an optrode that directly integrates the light source onto the recording electrode can significantly reduce technical complexity and provide a convenient way to conduct neuroscience experiments, thereby accelerating brain research.

Silicon was the first material used for attempts at incorporating an LED into an optrode due to the mature semiconductor processing technology. Growing Gallium Nitride (GaN) LEDs on silicon substrates, Wu et al. fabricated an optrode array with four prongs, each prong having 12 micro-LEDs and 36 recording sites^14^. However, Due to the photovoltaic effect in the silicon substrate, artifact noise for the recording electrodes could be quite high during LED illumination. To address this issue, Kim et al. fabricated a micro-LED optrode using heavily boron-doped silicon-based GaN ^15^ to reduce signal noise. Despite the success of combining silicon and GaN technology to create optrodes, there are a few shortcomings associated with this technology which make this approach less than ideal. For instance, silicon is rather brittle and is known to be easily breakable while being inserted into brain tissue. Additionally, a larger number of surface defects can be generated due to the significant lattice mismatch and thermal stress between GaN and the silicon substrate when GaN is grown on the silicon substrate^16^. This leads to lower light emission efficiency for the LED^12^. Silicon is also a non-transparent material, so light emitted from the LEDs can only optogenetically excite neurons on the same side as the LED. To achieve uniform illumination of the brain nuclei for better behavioral responses, LEDs would need to be grown on both sides of the silicon probe, which can be technologically challenging.

Sapphire is a strong material with a hardness second only to diamond, exceptional thermal conductivity for dissipating heat, and good transparency to light. Additionally, the emissivity of GaN LEDs grown on sapphire is higher than those on silicon substrates due to better lattice matching between GaN and sapphire^17^. Sapphire, therefore, has good potential as an alternative material for creating optrodes for neuroscience studies. McAlinden et al.^18^ created an optical probe with five independently addressable miniaturized GaN LEDs grown on sapphire and demonstrated that the LED probe could induce action potentials in-vivo^19^, but this probe lacked the capability to record action potential firing. After this work, Zhang et al.^20^ monolithically integrated a GaN LED on a sapphire shank with eight neural electrodes and measured the optical and electrical characteristics of their probes; however, no cell or animal recordings were done to support the functions of these optrodes. Recently, a 10×10 sapphire needle waveguide was also created to deliver GaN-LED generated light to stimulate deep cortical layers of non-human primates^21,22^. In our previous work, we have used sapphire as the optrode substrate^23^ and developed a double-sided design by growing an LED and recording electrodes separately on the top and bottom sides of the sapphire substrate separately^24^. Since the sapphire substrate is transparent, the LED light can transmit to the other side of the electrode for optogenetic stimulation. Although we have not yet performed experiments with sliced brain tissues or animal models to confirm the functions of the sapphire optrode, we have characterized the electrical and optical properties of this design.

A common issue for both silicon and sapphire optrodes in which recording electrodes and optogenetic LEDs were integrated in close proximity on the same shaft is noise crosstalk between the recording electrodes and the LED. This noise crosstalk adds additional noise to the recorded neural voltage, especially at the onset of the LED operation, which can overwhelm the neural signal. In this work, multiple layers of grounding structure were built into the optrode to electrically separate the recording sites from the LED to avoid this noise crosstalk. This electrical insulating effect was modeled and experimentally confirmed to reduce this noise crosstalk. In addition, metal electrodes are susceptible to photochemical effect and photovoltaic effect under light illumination, and the photovoltaic effect (PV) can also generate additional noise with the silicon substrate^25–27^. In this regard, sapphire has a much higher band gap (Eg=8.1 eV) than that of silicon and therefore provides better electrical insulation, making it naturally more resistant to this PV noise than silicon.

In this paper, we discuss further improvements to our sapphire optrode and demonstrate its simultaneous neural recording and optogenetic stimulation functions by measuring neural activities of the medial superior olive (MSO) in anesthetized Mongolian gerbils. The sapphire optrode integrates a GaN 480nm blue LED directly on top of a neural recording array with 10 (5×2) recording sites to ensure co-registering of the neurons for optogenetic stimulation and neural recording. Sapphire provides a rigid structure, allowing for the creation of a long probe (35 mm) with a high aspect ratio and minimal deflection to precisely target deep brain nuclei. A multilayer grounding structure is also incorporated within the optrode to shield the recording electrodes from the electrical artifacts induced by the onset of LED illumination. The sapphire optrode offers an easy-to-use package for studying neuronal circuits’ functions using both electrophysiology and optogenetic manipulation.

## RESULTS

### Design of the sapphire optrode

An optrode, capable of recording neural spike activities at multiple locations and simultaneously optogenetically stimulating neural activities, was designed and fabricated based on a sapphire substrate. Compared to typical silicon substrates used for making neural probes, sapphire has superior properties, including stronger mechanical strength and transparency for light transmission. In our current design, the sapphire optrode consists of 10 metal recording sites arranged in a 2×5 array for neural recording and a blue LED for channelrhodopsin-related neural stimulation. Fig. 1a and b show bird’s-eye and close-up views of the optrode with the 10 gold (Au) electrodes, each with a circular diameter of 30 µm, arranged in a 2×5 array with a site-to-site separation of 60 µm, on the insertion side (left in the figure). A 300×140 µm GaN blue LED with a peak emission wavelength of 458 nm was grown directly underneath the recording array. In this paper, a Dielectric Bragg Reflector (DBR) was grown beneath the sapphire substrate to reflect the LED light back to the front surface for more effective optogenetic stimulation. This DBR can optionally be removed to allow light to stimulate neurons on the back side of the optrode, due to the transparent nature of sapphire. Fig. 1c is an expanded cross-sectional view of the optrode showing all the layers constructing the optrode. Three ground interlayers (the 2nd, 4th, and 6th layers from the top) were used for noise shielding and co-served for other functional purposes. The Ti/Al/Ni/Au (Shielding-1) layer grown on top of the SiO₂ insulation and MQW layers covers a significant portion of the entire electrode for noise shielding, while also having separate power wires for the cathode and anode of the MQW LED to carry power to the LED. The second Au (Shielding-2) layer consists of a thick shielding line for noise shielding and ten interconnection wires to connect to the 10 recording sites for neural signal transmission. The top Cr/Ti (Shielding-3) ground layer covers the entire electrode to provide maximum noise shielding, with a large opening exposing the LED and the recording arrays. The width and thickness of the optrode are 490 µm and 150 µm, respectively, and the length of the optrode is 3.5 cm, which is long enough to reach deep nuclei, such as neural targets in the midbrain.

**Figure 1.**
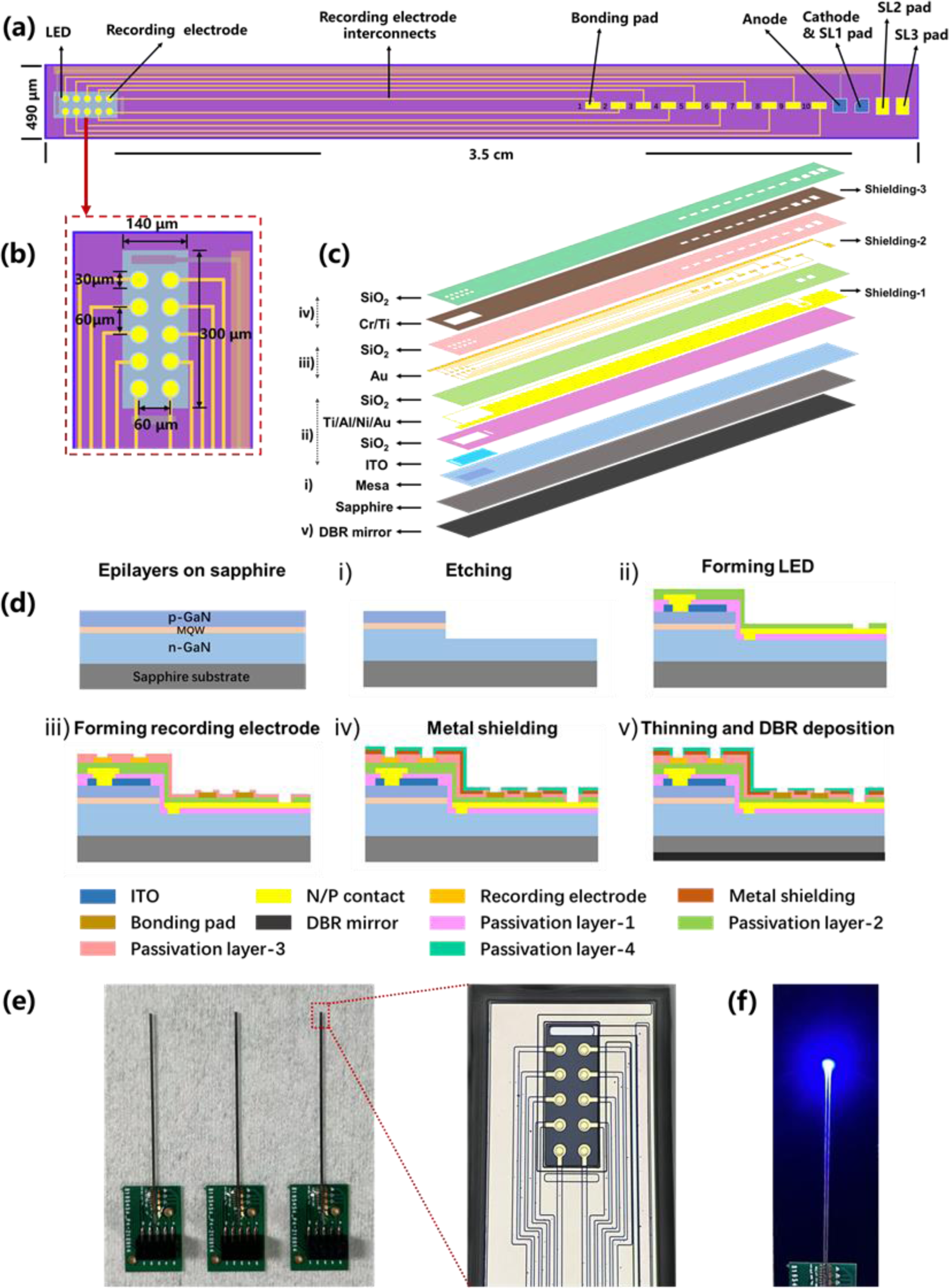
(a) Bird’s-eye schematic of the sapphire optrode; (b) Close-up of the LED and the recording electrode area of the optrode; (c) Expanded cross-sectional view of the optrode showing the layers; (d) Fabrication process of the sapphire optrode; (e) Photograph of three completed optrodes and a zoomed-in photograph of the tip of the optrode; (f) Optrode LED at work emitting 458 nm photons. Previously silicon has been used as the base material for most neural recording electrodes and optrodes, but sapphire is not yet commonly utilized for this application. For this reason, the mechanical, optical, electrical, and performance stability of our optrode were characterized for reference.

Fig. 1d illustrates the fabrication process of the sapphire optrode. A sapphire substrate grown with an MQW LED layer sandwiched between p– and n-type GaN layers was ultrasonically cleaned in acetone and isopropanol and then dipped into an acidic solution to remove inorganic impurities. A mesa with a height of 900 nm was then etched onto the GaN layer using Inductively Coupled Plasma (ICP) to form a blue LED. A 50 nm thick ITO thin film was deposited onto the layers using magnetron sputtering and annealed in nitrogen at 500 °C for 15 minutes to create ohmic contacts. A silicon dioxide (SiO₂) isolation layer was deposited using Plasma Enhanced Chemical Vapor Deposition (PECVD) to cover the LED. A layer of Ti/Al/Ni/Au metal was then vapor-deposited using electron beam evaporation onto the ITO layer and etched to form connections to the LED underneath. This metal layer also served as the first ground layer (Shielding-1), shielding the LED from electric noise. After the LED was made, another SiO₂ layer was deposited as a passivation layer to isolate the LED from the recording sites to be created. A layer of gold (Au) metal was deposited above the SiO₂ layer using electron beam evaporation, followed by etching to form the 2×5 neural recording arrays. This Au metal layer also served as the second shielding layer by embedding a long and thick ground wire (Shielding-2) along the optrode body. A third SiO₂ isolation film was then deposited covering the recording signal wires but also etched to expose the recording sites. A Cr/Ti metal layer was deposited covering the entire electrode to serve as the third ground shielding layer (Shielding-3), except a square was etched to expose the recording sites. The top of the electrode was covered with an SiO₂ isolation film to protect the electrode. Once the optrode substrate is thinned and polished, a DBR mirror layer can be optionally grown on the back of the optrode to reflect the back-emitted LED light to the front. Lastly, the sapphire wafer was isolated into individual optrodes with laser cutting.

After the fabrication of the optrodes, the output ends of the optrodes were glued onto a printed circuit board (PCB) and the conductive pads of the optrode were ultrasonically wire-bonded to the connective pads of the PCB with aluminum wiring, allowing the optrode to be connected to pre-amps and other external electronics. The wire bonding area of the PCB was covered with photosensitive resin to provide insulation and mechanical protection for the metal bonding wires. The weight of each optrode was measured to be 0.33 g. Photographic images of encapsulated optrodes and a zoomed-in photograph showing the recording sites and the LED are shown in Fig. 1e. Fig. 1f also shows the sapphire electrode with the blue LED turned on emitting 458 nm photons.

Previously silicon has been used as the base material for most neural recording electrodes and optrodes, but sapphire is not yet commonly utilized for this application. For this reason, the mechanical, optical, electrical, and performance stability of our optrode were characterized for reference.

### Mechanical properties of the optrode

Fig. 2a shows the insertion displacement of the sapphire optrode when a hard tip was pressed onto its top surface with a load ranging from 0 to 1 N. The measured displacement curve exhibits hysteresis with a maximum indentation of 2725.6 nm. Using the measured displacements, the modulus and hardness of the optrode were estimated and are plotted in Fig. 2b and c. As the tip pressed onto the optrode surface, the elastic modulus of the optrode initially decreased over the first 250 nm and then increased linearly, reaching a maximum of 133.29 GPa. Similarly, the hardness of the optrode increased rapidly at first and then plateaued around 8.5 GPa, reaching 10 GPa at maximum displacement. This initial nonlinear behavior was likely due to the optrode’s non-uniform construction, consisting of multiple layers of various softer materials. As the tip pressed deeper into the material, these different layers compounded, causing variations in the mechanical properties of the optrode. The impression formed by the hard tip was imaged with an Atomic Force Microscope (AFM) (Fig. 2d). Additional AFM photos are available in the supplementary information.

**Figure 2.**
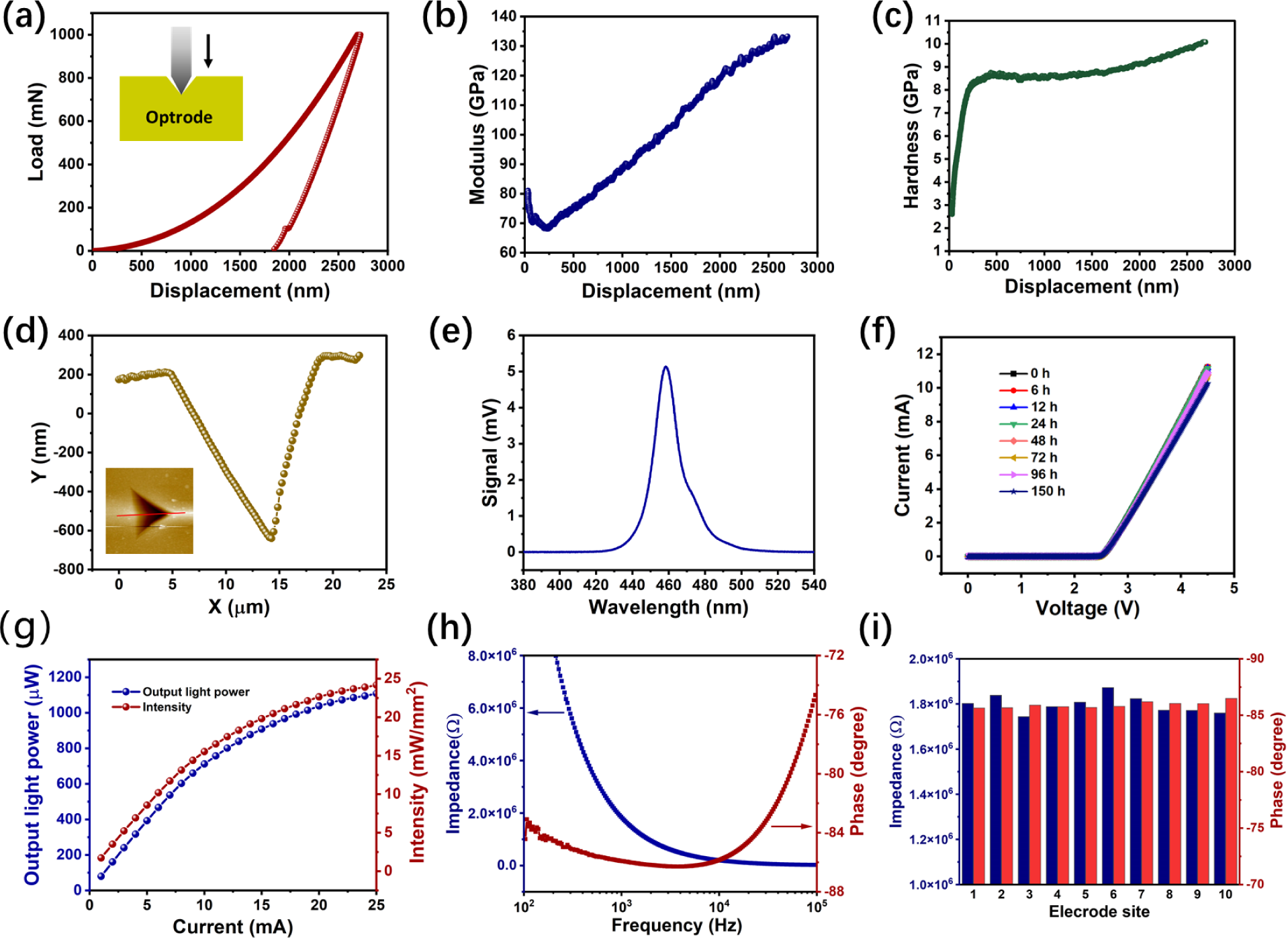
(a) Insertion displacement load of the sapphire optrode; (b) Modulus and (c) hardness of the sapphire optrode against insertion displacement; (d) Impression formed by the hard tip insertion and the corresponding AFM image of the optrode surface; (e) Photoluminescence Spectrum (PL) of GaN LED; (f) I-V curves of the optrode LED after soaking in PBS for 0 to 150 h; (g) LED optical power and intensity against driving current; (h) Averaged amplitude and phase of the electrode impedance; (i) impedances and phases measured at 1 kHz for each of the 10 electrodes.

### Optical properties of the optrode

Fig. 2e shows the measured photoluminescence (PL) spectrum of the optrode LED. The LED fluoresces from 440 to 480 nm, peaking at 458 nm, which matches well with the absorption wavelengths of most excitatory opsins used for optogenetic stimulation. The current-voltage (I-V) curve of the LED was measured, and the LED turned on at 2.3 V (threshold voltage) and could be driven as high as 4.5 V without observable degradation. Since most optogenetic experiments require the optrode to be implanted in animal brains for extended periods, this situation was simulated by submerging the optrode in phosphate buffer solution (PBS, 0.01 mol/L, pH 7.2-7.4) solution for up to 150 hours. The I-V curves measured after the extended submersion do not show any significant differences compared to the curves measured before submersion (Fig. 2f), indicating that the LED can be continuously used in an aqueous environment without affecting its light emission capability. The optical power output of the optrode LED was measured against the input current (Fig. 2g). The optical power increased linearly in the at lower currents, (0 to 15 mA) but slightly saturated at higher driving currents (15 to 25 mA), likely due to heating effects. A maximum optical power of 1030 µW was measured at a driving current of 25 mA, corresponding to a maximum optical intensity of 24.6 mW/mm². This optical power greatly exceeds the activation optical intensity required for most optogenetic opsins.

### Electrochemical characterization of the recording electrode

The electrical impedance of a neural recording electrode is often used as a parameter to characterize its neural recording performance. The impedance of the sapphire optrode was measured from 100 to 10,000 Hz, and the amplitude and phase of the impedance are shown in Fig. 2h. The impedance of the optrode decreased exponentially with frequency, likely due to the reduction of the effective capacitance at the interface between the metal recording sites and the PBS solution. Specifically, the amplitude and phase of the impedance at 1 kHz were measured to be 1.8 MΩ and –86°, respectively. Additionally, the impedances of 10 recording channels on the optrode were measured to determine variations between the electrodes. Fig. 2i shows that the impedance and phase angle variations measured at 1 kHz over the ten recording channels were less than 1%.

### Multi metal shielding effectively reduce EMI-induced artifact

The sapphire optrode described in this manuscript successfully integrates a blue LED directly underneath recording channels to allow simultaneous neural recording and optogenetic stimulation in an easy-to-use package. However, the close proximity between the recording sites and the LED can cause significant electromagnetic interference (EMI) from the LED to be picked up by the recording channels. To mitigate this issue, ground shielding layers were embedded within the optrode to electrically insulate the LED from the recording sites. Four shielding scenarios were simulated in 2D – full shielding with ground, full shielding with no ground, and no shielding – to investigate the strength of the shielding effects. The simulated false color voltage plots are shown in Fig. 3. For the simulation, the LED was set to have a constant voltage potential of 3 V and the three shielding layers were either grounded (voltage set to 0 V), left floated, or completely removed. The first scenario (all shielding layers present, voltage set to 0 V) is illustrated in Fig. 3a. Due to the grounding, the voltage potential within the optrode was estimated to be close to 0 V, essentially suppressing the voltage potential emanating from the LED (Fig. 3a right). To understand the effects of each grounding layer, the top and middle interlayers were still present but left potentially floated in the second scenario. Without grounding these two layers, the potential near the recording electrodes rose to ∼0.5 V (Fig. 3b), indicating that the grounding of the top layers is important in preventing stray potential from reaching the recording sites. If the top two interlayers were completely removed from the optrode structure, the voltage potential near the recording electrodes further increased from ∼0.5 V to ∼0.7 V (Fig. 3c). If the bottom shielding layer was also removed from the optrode, the voltage potentials at the recording sites were as high as ∼2.2V. These simulated results indicated that the embedded ground layers can be very effective in shielding the recording electrodes from EMI. Although this simulation was performed only with static electromagnetic simulation, these layers should also be effective during voltage onsets, such as turning on and off the LED, thereby reducing the electromagnetic distortion caused by the proximity of the LED to the recording sites.

**Figure 3.**
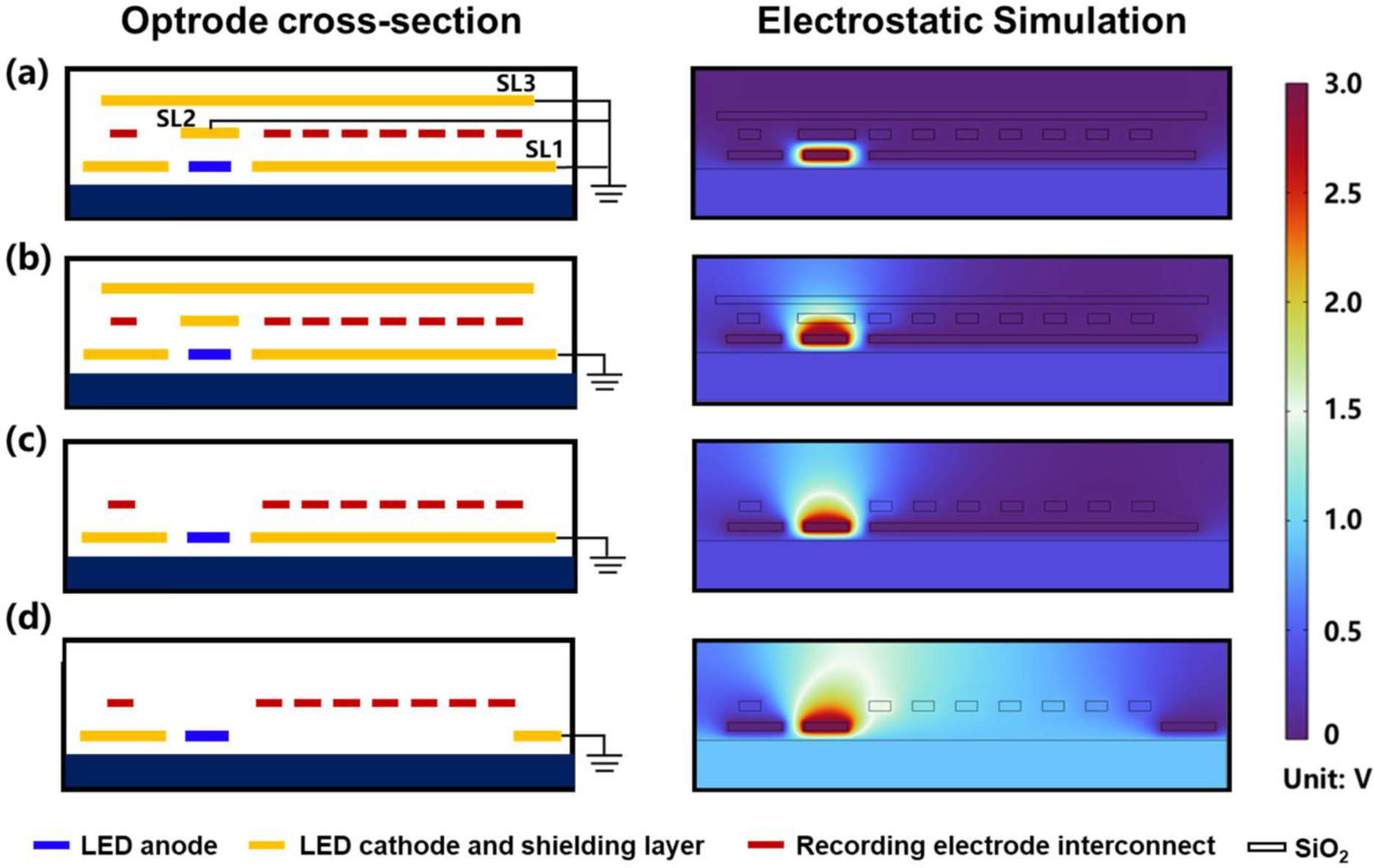
2D structures (left) and electromagnetic simulations (right) of the sapphire optrode incorporating three shielding interlayers under different grounding conditions. (a) full ground shielding with all 3 interlayers (b) ground floating for the top two interlayers; (c) top shielding interlayer removed; and (d) without ground shielding.

Controlled experiments were also conducted to compare the recorded voltages with and without grounding the embedded shielding layers. Fig. 4a illustrates the experimental setup used to compare the recorded voltages. A sapphire optrode was dipped into a glass beaker filled with PBS to mimic an optrode submerged in the brain. The optrode was then attached to a custom-built multi-channel bio-amplifier and an analog-to-digital data acquisition board for recording. The bio-amplifier was powered by a battery pack to avoid introducing power line noise onto the recorded voltage signals. A pulse generator was connected to the optrode LED to drive light emission with repetitive square pulses (3.0 V, 10 Hz, and 50% duty cycle), and voltages at the recording sites induced by the optrode LED’s electrical dynamics were measured. A common ground was used to connect all instrumentation, as well as the optrode and the PBS solution, during the test. The shielding interlayers were subjected to different grounding conditions, either floated or grounded, and the measured results are shown in Fig. 4b. When the metal shielding layer was absent, the measured peak voltage could reach a maximum amplitude of ∼7.0 mV, but the voltage of a fully shielded and grounded optrode had a measured peak amplitude of only 4.5 mV—a reduction of more than 40%. Besides the peak height difference, the total voltage areas of the unshielded and shielded cases were also different. For the unshielded case, the peak voltage of the unshielded optrode did not immediately decay but first plateaued at 7.0 mV for approximately 2 ms before it exponentially reduced to the baseline level. In contrast, for the shielded case, the measured voltage of the shielded optrode immediately reduced after reaching its maximum value. These differences indicate that the shielding layers are effective in reducing, albeit not eliminating, the impact of EMI generated by the emission onset of the LED.

**Figure 4.**
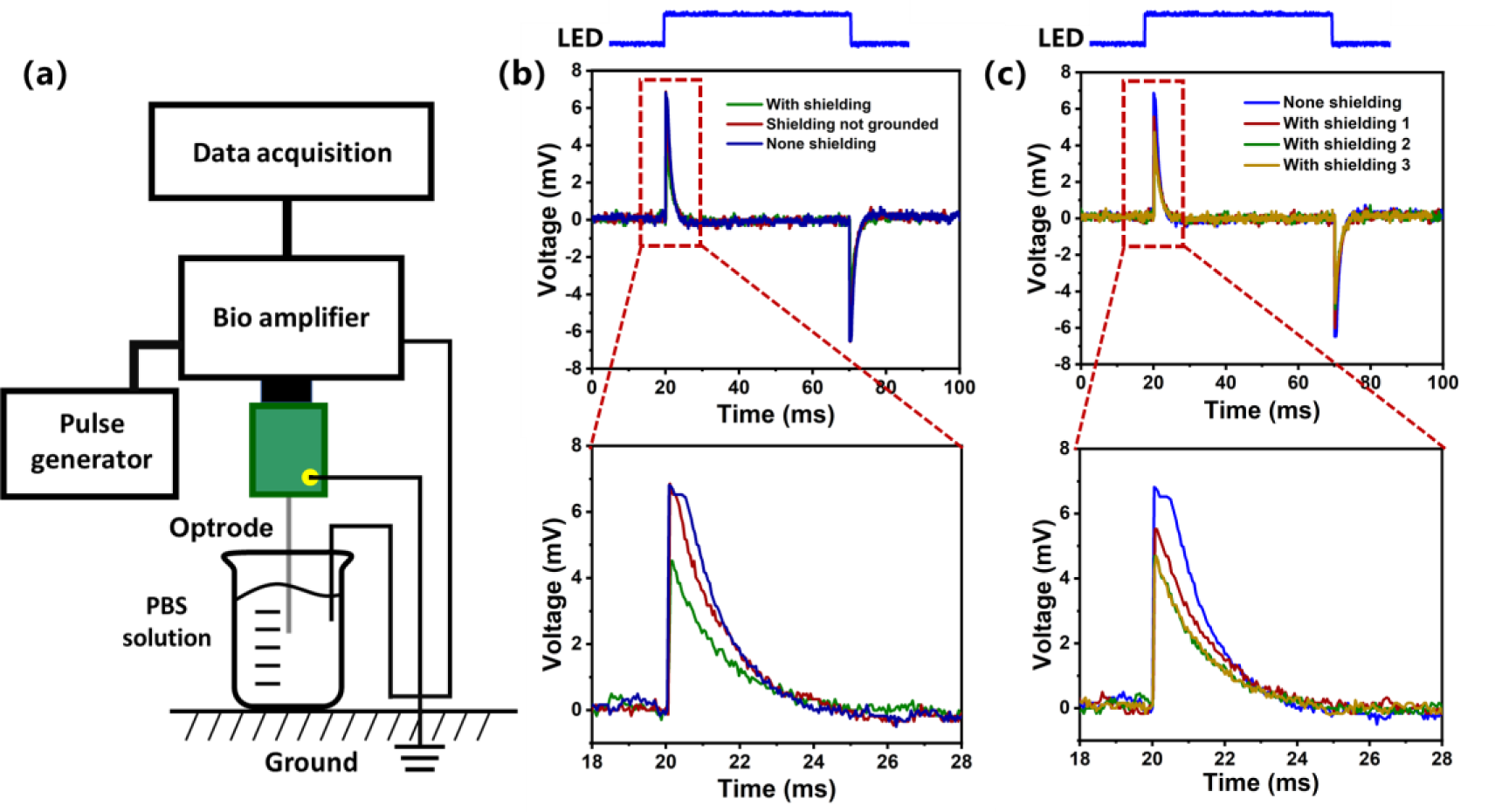
(a) In-vitro experimental schematic evaluating LED induced noise on the recording electrodes; (b) Measured voltage time traces when the optrode interlayers were fully grounded, shielded but not grounded, and not shielded during the onsets of the optrode LED; (c) Measured voltage time traces when the shielding interlayers are not shielded and with shielding layers.

### In-vivo extracellular recording using the sapphire optrode

The neural recording capability of the optrodes was demonstrated by recording mitral/tuft cells from the olfactory bulb of a C57BL/6 mouse under anesthesia. The schematic of the experimental setup is illustrated in Fig. 5a, and the recorded neural voltage trace is plotted in Fig. 5b. The root mean square value of the recorded trace was measured to be 18.225 ± 4.48 µV_rms_ (mean ± standard deviation; 100 1-second segments). This noise level is comparable to that of tungsten electrodes with similar impedance. Neural spikes of the mitral/tuft cells measured by the sapphire optrode had a good signal-to-noise ratio, reaching an average SNR of 13.61 ± 1.83 dB_rms_ (mean ± standard deviation; 100 1-second segments), i.e., neural spike signals are approximately 4.79 to 5.91 times higher than the noise floor and could be easily separated in following analysis. After recording, the measured neural spikes were isolated and spike-sorted to reveal single-unit activities. Three single-unit neural spike shapes were identified, as shown in Fig. 5c. These spike clusters had distinct temporal profiles and good conformity to the profiles, indicating that the electrophysiology recording capability of the sapphire optrodes is reasonably good for measuring neural activities in-vivo.

**Figure 5.**
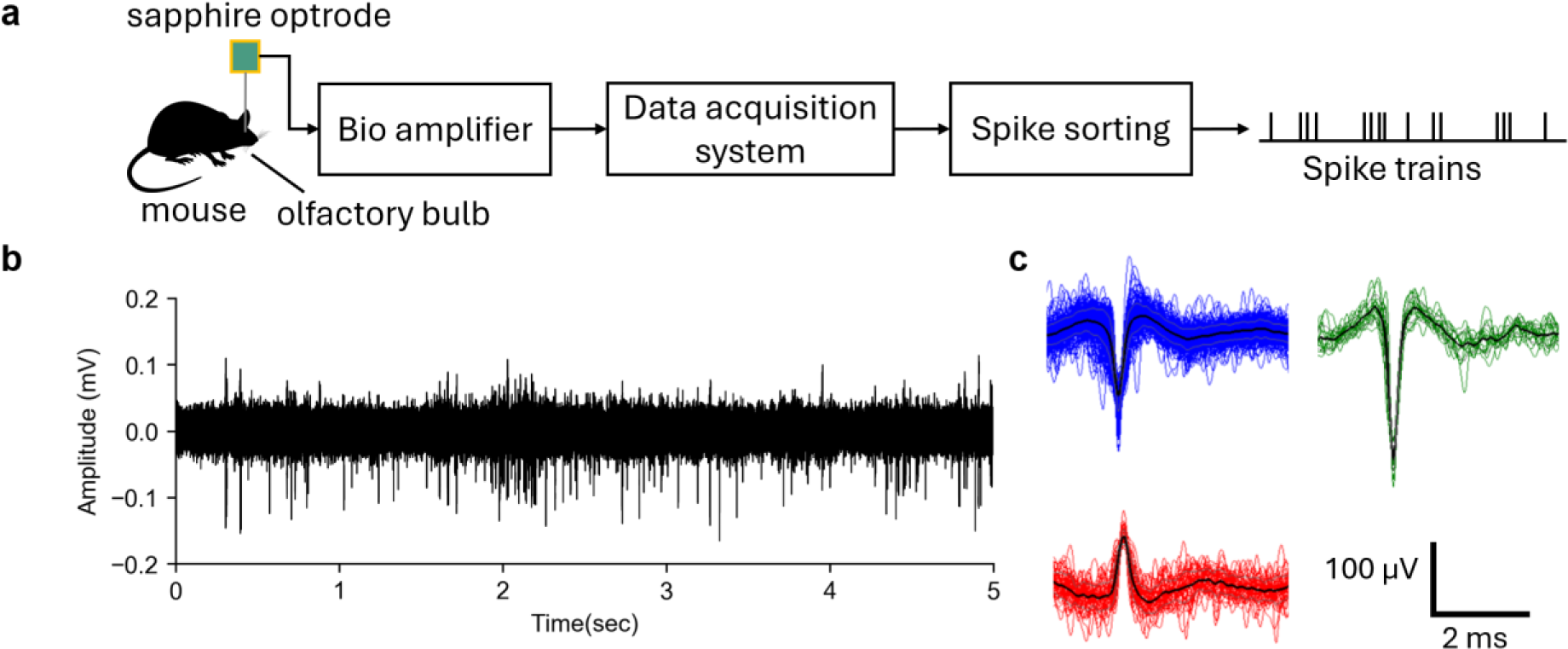
In-vivo extracellular recording using the sapphire optrode. (a) The experimental schematic for recording neural action potentials from mitral/tufted cells of the olfactory bulb of an anesthetized mouse; (b) recorded neural voltage showing action potentials embedded on a noise background with an average SNR ratio of 13.61 ± 1.83 dB_rms_ (mean ± standard deviation; 100 1-second segments); (c) isolated neural spikes after analysis by a spike sorting routine, showing these neural spikes originating from three different neurons.

### In-vivo optogenetic manipulation using sapphire optrode

The sapphire optrode, which monolithically integrates an LED with recording electrodes, provides a convenient experimental package for simultaneous neuronal activity measurement and optogenetic neural manipulation. In this section, neuronal firing activities of opsin-expressing medial superior olive (MSO) neurons were recorded using the sapphire optrode while optical stimulation was delivered during an auditory experiment, as illustrated in Fig. 6a. The location of the left MSO nucleus for the viral vector injection and in-vivo extracellular optogenetic recording is illustrated in Fig.6b. The MSO is known to be an important auditory processing unit in multiple aspects of hearing and is a crucial component of the ascending and descending auditory pathways; therefore, it is electrophysiologically responsive to sound stimulation ^28^. After a 3-week incubation period after injection, the Channelrhodopsin-type (ChR2) opsin was expressed in the MSO region, which was confirmed by anatomical verification as shown in the confocal image in Fig.6c. An example neural trace recorded under repetitive 50 ms sound stimulation (auditory-evoked response) is displayed in Fig.6d, where the LED of the optrode was turned on (indicated by the blue line at the top) to stimulate the opsins expressed in the MSO neurons. The example trace demonstrates stronger auditory-evoked responses under light stimulation.

**Figure 6.**
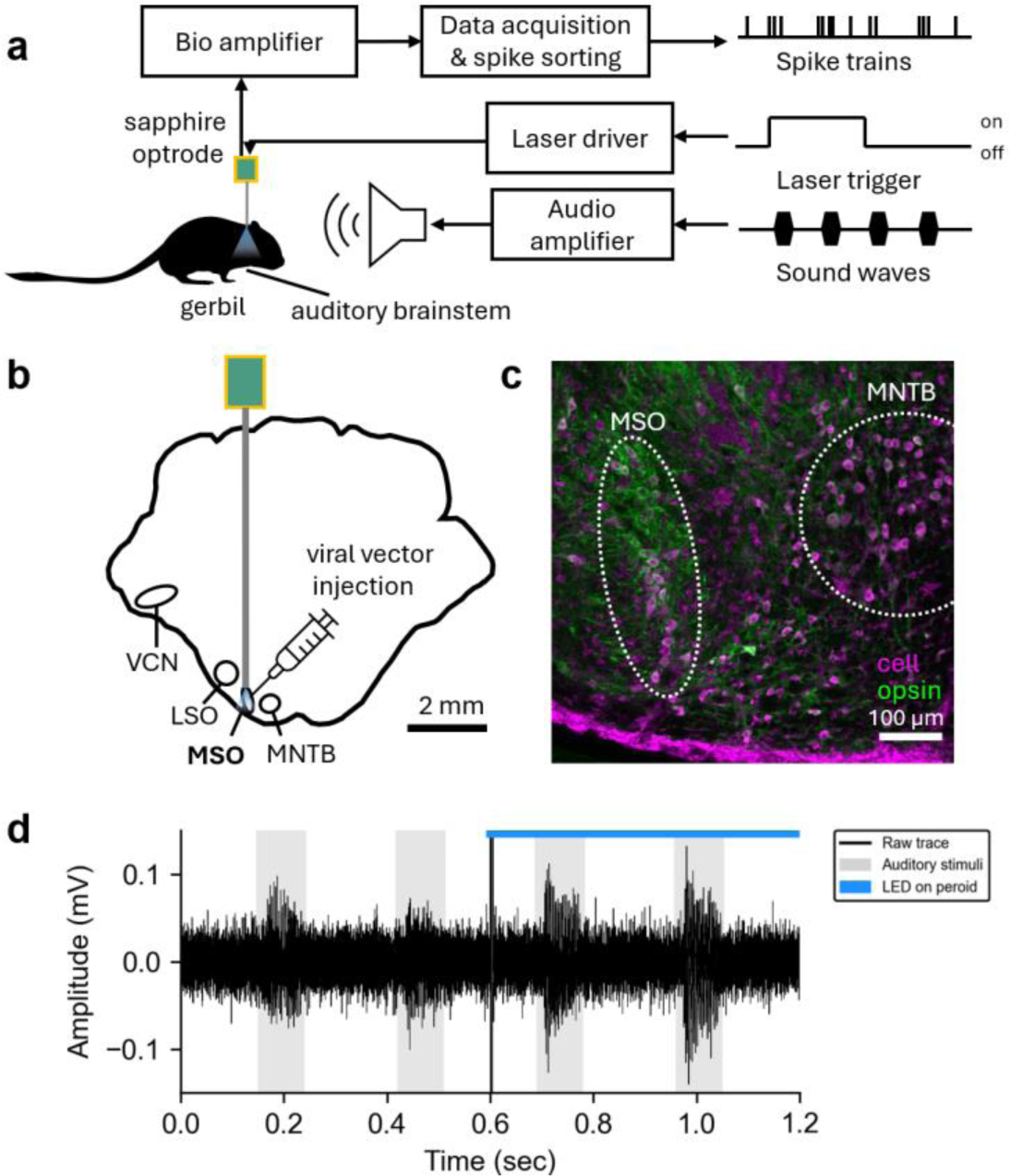
In-vivo optogenetic recording using a sapphire optrode to record and optogenetically stimulate auditory brainstem neurons. (a) A schematic of the optogenetic experiment, where medial superior olive (MSO) neurons in the auditory brainstem of Mongolian gerbils were recorded while sound stimuli and optical stimuli were presented. The targeted brain region was injected with channelrhodopsin-based viral vectors at least 3 weeks prior to the recording experiment. (b) Simplified coronal section of the gerbil auditory brainstem indicating the location of MSO and related nuclei (LSO: lateral superior olive; MNTB: medial nucleus of the trapezoid body; VCN: ventral cochlear nucleus). (c) Confocal microscopy image of the targeted brain area expressing opsins through viral vector injection. (magenta color: cells; green color: opsin expression; scale bar: 100 micrometers) (d) A typical raw data trace recorded from MSO. The noise floor did not differ under light stimuli except a short onset transient (blue triangle), while the auditory-evoked responses become more severe under light stimuli. (Black line: raw trace; gray shading: sound stimuli presented; blue bar: LED light presented; blue triangle: LED onset time)

Peri-stimulus time histograms (PSTH) of the recorded MSO neural spikes are shown when the LED was turned off (Fig. 7a) and turned on (Fig. 7b) under repetitive pure tone stimulation (100, 125, 150, 175, and 200 Hz). During pure tone stimulations (grey areas), MSO neural firing rates increased to varying degrees depending on the pure tone frequencies, indicating that the sapphire optrode was targeting MSO neurons tuned to roughly 150 Hz sound stimulation. Comparing the firing rates when the LED was turned on and off (Fig. 7c and d) showed that the firing rates significantly increased when the LED was turned on. Since the only variable between the two scenarios was the activation of the LED on the sapphire optrode, the increased firing rates can only be attributed to the successful activation of the opsins expressed in the MSO neurons. Additionally, when the sound pressure level (SPL) of the two speakers was increased from 20 to 80 dB, the firing rates increased at roughly the same rate for both scenarios. However, the firing rates of the MSO neurons were significantly higher when the LED was turned on, as shown in Fig. 7e. The Interaural Time Difference (ITD), in which the onsets of the pure tones on the left and right ears were temporally offset, was also measured. The results show the typical ITD tuning response for MSO neurons (Fig. 7f). The tuning curve when the LED was on again shows an increased firing rate, indicating that the LED on the sapphire optrode successfully activated opsins expressed in the MSO neurons. Finally, it is noteworthy that the onset of the LED elicited a brief transient artifact pulse but did not impact the subsequent noise floor.

**Figure 7.**
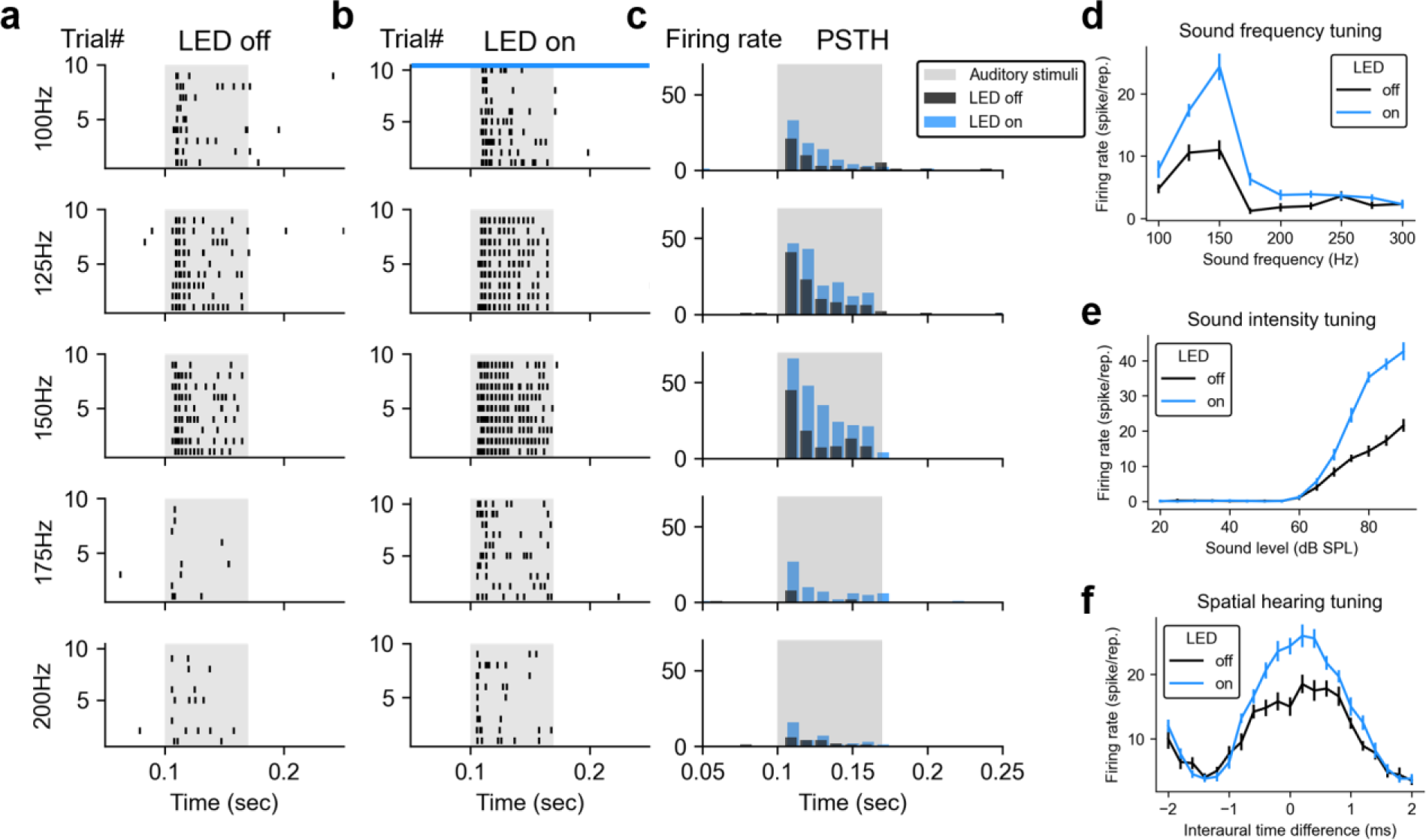
Modulated auditory-evoked neural responses in the in-vivo optogenetic experiment using proposed optrode. Raster plots of single-unit neural spikes from the example MSO neuron with (b) and without (a) light stimuli presented across 10 trials of repeated auditory stimuli with different frequencies. (Black dots: neural spikes; gray shading: sound stimuli presented; blue bar: LED light presented) (c) Peri-stimulus time histograms (PSTH) calculated from the example MSO neuron under different sound frequencies. (Black bars: neural firing rates without light stimulus; blue bars: neural firing rates with light stimulus presented; gray shading: sound stimuli presented) (d) Neural firing rates of the example MSO neuron under different sound frequencies with (blue line) and without (black line) light stimulus presented. (Error bar: standard error) Neural firing rates of the example MSO neuron under different sound intensities (e), and interaural time differences (f, a kind of binaural cue for the sound localization process indicating the arrival time difference of sound between two ears) with (blue line) and without (black line) light stimulus presented. (Error bars: standard error)

## DISCUSSION

In this paper, we report on the design, manufacturing, and evaluation of our sapphire optrode that monolithically integrates a blue LED and 10 neural recording sites for simultaneous electrophysiological recording. The sapphire optrode has been successfully demonstrated to measure action potential firing from the olfactory bulb and optogenetically elevate action potential firing in the MSO of anesthetized Mongolian gerbils. This optrode provides an easy-to-use package for conducting optogenetically driven electrophysiology measurements in-vivo and potentially in behavioral neuroscience studies.

Despite the existence of silicon optrodes, sapphire optrodes offer several advantages (Table 1). Sapphire is mechanically much more rigid than silicon, and silicon probes or optrodes are known to break easily during insertion into animal brain tissues. On the other hand, sapphire optrodes are much more robust and can be reused multiple times by cleaning them with isopropanol. From a material perspective, GaN-on-sapphire has an order of magnitude lower dislocation density (3×10⁸ cm⁻²) compared to GaN-on-silicon (5×10⁹ cm⁻²). Consequently, the light emission efficiency of GaN LEDs grown on sapphire substrates is much higher. Due to this increased emission efficiency, under the same driving current, sapphire optrodes can produce more light to excite deeper neurons, better covering the entire perimeter of the target neuron optically. Additionally, silicon-based probes tend to generate photovoltaic noise when illuminated by light, primarily originating from the silicon substrate underneath, which can adversely affect the measured neural signals^29^. Some researchers have tried using heavily doped silicon substrates or flexible substrates to mitigate this issue^12,27,30,31^. However, since the bandgap of the sapphire substrate is much higher than that of silicon, this photovoltaic noise effect is naturally much reduced. Additionally, the transparent nature of sapphire allows for more comprehensive light coverage of the targeted nuclei, providing a much better behavioral response in manipulating neuronal circuits for cleaner experimental outcomes.

**Table 1.**
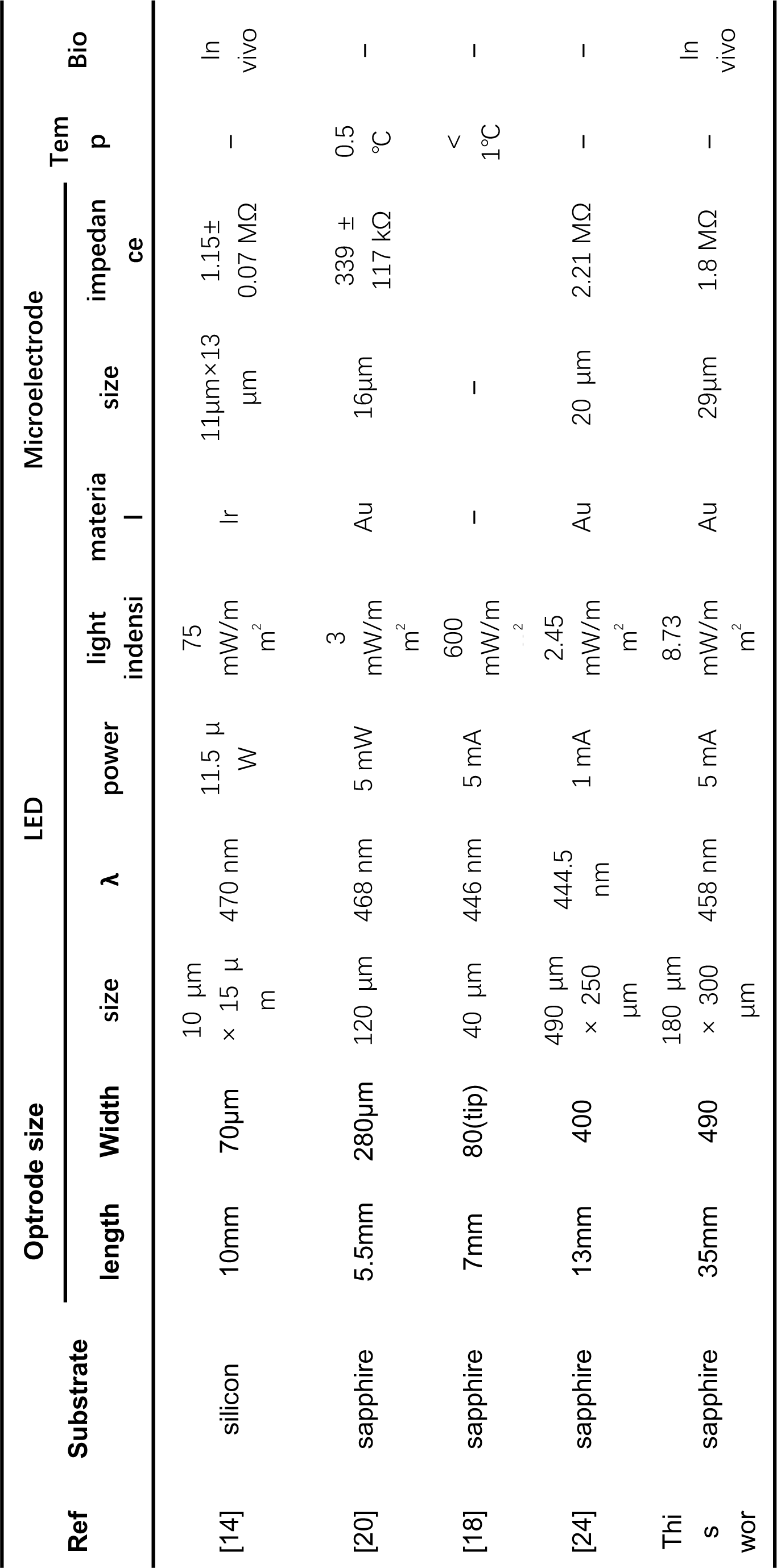
Comparison of performances and parameters for rigid LED optrode.

For the purposes of demonstrating the ability of the optrode to simultaneously record and optogenetically modulate neural activity, we targeted the MSO. Instead of targeting a shallower neural target, MSO neurons were chosen because they are challenging to record from and optogenetically manipulate. Anatomically, the MSO is a tiny nucleus located near the ventral side of the brainstem, with neurons arranged in almost a single cell layer extending dorsoventrally (Fig. 6 b and c). Therefore, targeting such a deep nucleus requires an optrode that is long and rigid enough to travel approximately 6-8 mm from the skull surface with minimal deformation. In this regard, sapphire is an excellent material for making long electrodes due to its strong material properties. Additionally, the MSO is close to several large fiber bundles traveling through the area of the superior olive and trapezoid body, making MSO neuron signals notoriously difficult to record ^32^. Optogenetically, this brain area has a very high optical scattering coefficient due to the heavy myelination of the passing fibers, making light penetration for neural stimulation difficult ^33^. Despite these challenges, the success in elevating action potential firing in the MSO indicates that the integrated LED can generate adequate light intensity to activate the opsins expressed in MSO neurons. This suggests that the optrode should be adequate to optogenetically stimulate most neural targets in the brain.

As mentioned in the introduction, there have been previous efforts to use sapphire as an optrode substrate. Table 1 lists selected silicon and sapphire neural probes ^14,18,20,24^ and compares them to ours, detailing their dimensions, number of LEDs, number of microelectrodes, impedances, and experimental demonstrations. Our sapphire probe is the only one that has been validated in animal models by simultaneous neural recording and optogenetic neural stimulation. Additionally, our neural probe incorporates a multilayer shielding structure to reduce electric noise artifacts during the onset of LED operations. DBR reflectors were incorporated on the back of the sapphire substrate to help redirect light to the neurons on the electrode side, improving optogenetic stimulation efficiency.

Despite the described sapphire optrodes having achieved our initial goal of successfully optogenetically stimulating neural activities and simultaneously recording action potentials, there are several shortcomings that should be further explored. In the current design, multiple shielding layers have been incorporated into the optrode to suppress noise; however, this noise was not fully eliminated. There are several possible explanations: shielding layer 2 was grown on the same layer as the microelectrodes, which could negatively impact the anode current and prevent optimal shielding. Despite grounding the shielding layers, there might still be parasitic capacitances or inductances between them, causing electromagnetic interference. Additionally, the microelectrodes could produce photoelectrochemical responses under light illumination, causing photoelectric noise. These issues can be further optimized in future designs of the shielding layer structure. For instance, a Faraday shielding structure could be incorporated for the electrodes to further remove electromagnetic interference. To address the potential light-induced photoelectric noise, electrochemical processing on the electrodes can be adapted to modify the microelectrode surfaces, thereby reducing this noise and increasing the SNR of the measured action potentials. Furthermore, the shank of the optrode was intentionally designed to be relatively long to reach deep nuclei, but this length can make it susceptible to electromagnetic interference. This shortcoming can be mitigated by applying a conductive paint on the surface of the shaft to allow grounding during measurements.

Additional improvements could include increasing the integration density of the electrode sites and reducing the size of LEDs without sacrificing the performance of the neural probe. For instance, the number of LEDs and recording sites can be increased and distributed along the sapphire shaft in the longitudinal direction, enabling the design of more complex neuroscience studies, such as targeting and optogenetically manipulating multiple neural targets simultaneously. By leveraging the robust physical properties and stiffness of the material, targeting more challenging neural targets might become possible. The high customizability of fabricating the sapphire optrode also suggests the potential to target multiple nuclei by growing LEDs and distributing the recording sites along the shaft of the optrode. The distributed LEDs should be independently controllable, allowing optogenetic stimulation across multiple brain nuclei at well-defined temporal moments. Efforts can also be made to reduce the width and thickness of the sapphire optrode to minimize brain damage during insertion. The width of the optrode is currently limited by the width of the transmission lines required to span from the recording sites to the bonding pads. However, integrated circuit technology can be employed to help reduce the width of these transmission lines, potentially down to the sub-μm level. Additionally, since inhibitory optogenetic opsins require optical photons in the 650 to 700 nm range (orange color), integrating LEDs with different emission spectra should be investigated in future iterations of this design. Furthermore, the development of flexible versions of this optrode may reduce biological tissue mechanical property mismatch with the optrode, thus reducing inflammatory responses for long-term implantation. In this regard, the sapphire optrode may potentially be transferred to a non-rigid underlying substrate through substrate peeling technology to make the optrode much more flexible^34^. Inflammatory reactions can significantly affect the voltage sensing capability of the recording sites. Therefore, modifying the metal recording sites to incorporate slow-release medicine pores could make the electrodes more resistant to biomaterial depositions. For instance, anti-inflammatory agents could be deposited onto the recording metal surfaces through electrospinning, followed by encapsulating the drug with slow-release pores, prolonging the recording time of the optrodes^35–38^.

The current work demonstrates that our sapphire optrodes can record and optogenetically stimulate neural activities in animal models and that the integrated shielding ground interlayers can effectively suppress stimulation artifacts caused by the close proximity of the LED and the recording sites. Because of these unique capabilities, the sapphire optrode is an easy-to-use device optimized for electrophysiology and optogenetic animal experiments.

## MATERIALS AND METHODS

### Mechanical, optical, electrical stability and noise characterization of the optrode

The mechanical hardness of the sapphire optrodes was measured using a Nano Indenter G200X (KLA Instruments) with a hardened tip puncturing into the sapphire substrate. The pressure data was then used to estimate the modulus and hardness of the sapphire substrate. A Bruker Atomic Force Microscope (AFM) operating in tapping mode was used to image and measure the vertical profile of the impression. The photoluminescence spectrum of the optrode LED was measured using a PL spectrometer (Nanometrics RPM Blue) and the optical intensity of the LED was measured using an optical power meter (Newport 1919-R). The current-voltage (I-V) curves of the LED were also measured using a semiconductor device parametric analyzer (Agilent B1500A, Keysight). The electrical stability of the optrodes was tested by soaking the optrodes in phosphate buffer solution (PBS, 0.01 mol/L, pH 7.2-7.4) at room temperature for various periods of time. The I-V curves of the LED were then measured before and after soaking in the PBS solution for comparison. To measure the electrical noise fluctuations of the recording electrodes, an Apollo II high-performance neural recording system (Bio-Signal Technologies) connected to an oscilloscope (DSOX4024A, Keysight) was used for signal amplification and measurement.

### Electrochemical measurements

The impedance of the recording electrodes was measured by an LCR meter (E4980A, Keysight) driven with a 50 mV AC source. In addition, cyclic voltammetry was measured with an electrochemical workstation (CS300, CorrTest) in constant current mode using the three-probe measurement geometry in which an Ag/AgCl wire and a Pt wire were used as the reference electrode and the counter electrode, respectively. The optrodes were first submerged in a buffer solution for 10 minutes to allow for stabilization before measurement. They were then dipped in a PBS solution during the measurement.

### Electrostatic field simulation

Electrostatic simulation using COMSOL Multiphysics simulation package (COMSOL Inc.) was performed to estimate whether the embedded shielding layers can reduce electrical noise at the recording sites caused by the voltage potentials at the integrated LED. Three different shielding models – 1) full shielding with ground, 2) full shielding with no ground, and 3) no shielding – were created to estimate the electrical shielding effects. The models were modeled in the two-dimensional space either using silicon dioxide or empty space as insulation layers between the shielding layers. A voltage potential of 3 V was applied to the LED while the electrodes were electrically floated for the simulation and were measured to estimate the noise shielding efficacies of the shielding layers.

### In-vivo extracellular neural recording

The in-vivo extracellular neural recording experiments were performed at the University of Macau and the mouse procedures were approved by the Institutional Animal Care and Use Committees (IACUC) of the University of Macau. Female C57BL/6 mice (6-8 weeks old) were anesthetized with isoflurane and a craniotomy was performed above the right olfactory bulb at 4 mm anterior to Bregma and 1 mm lateral to the midline. The sapphire optrodes were connected to a custom-made bio-amplifier board and were mounted onto the stereotaxic platform for brain nucleus targeting and shielded by a Faraday cage for reducing environmental noise contamination. The tip of the sapphire optrode was then lowered to a depth of 1.9 mm from the surface of the skull to reach the ventral mitral cell layer ^39^ for recording. A needle was inserted subcutaneously at the dorsal neck as the ground reference. The recorded neural voltage amplified by the bio-amplifier was digitized by a data acquisition card (NIDAQ, National Instruments, TX) at a sampling rate of 125 kHz. Neural spikes isolated from the recorded neural voltage with a voltage threshold above the noise floor contained action potentials fired from several mitral neurons, and the single-unit activities were sorted with the Wave_Clus spike sorting algorithm^40^.

### Simultaneous extracellular recording and optogenetic manipulation

All experimental procedures were performed at the University of Colorado Anschutz Medical Campus in compliance with all applicable laws and guidelines from the National Institutes of Health and were approved by the University of Colorado Institutional Animal Care and Use Committee. The optogenetic experiments were conducted on Mongolian gerbils (Meriones unguiculatus), a rodent species with sensitive low-frequency hearing^41^. Gerbils are a commonly used model species in auditory physiological research due to the similarity between their hearing range and that of humans. The objective was to use the proposed sapphire optrode to optogenetically manipulate neural activity and test the effects of the manipulation on the sound localization process. Recordings were performed from medial superior olive (MSO) neurons in the auditory brainstem, which detect differences in sound arrival times between the two ears, i.e., interaural time differences^42^. Channelrhodopsin-type (ChR2) adeno-associated viral constructs expressing the excitatory opsin ChETA under a human synapsin promoter were stereotactically injected into the left MSO under isoflurane anesthesia using a nanoliter injector (WPI, Sarasota, FL) and a glass micropipette inserted at an angle of 20 degrees toward the rostral at a depth of 7500 µm. This was accomplished through a craniotomy made with a dental drill 4 mm caudal and 0.8 mm left of the lambda landmark. A total volume of 483 nL of the virus was injected through 15 injections, with one injection administered every 15 seconds. The animals were allowed to recover for at least three weeks before any further experimentation was performed to allow for viral expression of the transgene.

During the subsequent recording session (Fig. 6a), gerbils were anesthetized with isoflurane and a second craniotomy was performed as described above. A needle electrode was inserted into the muscle of the dorsal neck as the ground reference. After surgery, isoflurane administration was terminated, and the animals were kept in the surgical anesthetic plane with intraperitoneal injections of a ketamine/xylazine mixture (induction dose: 5 mg ketamine/100 g BW and 0.25 mg xylazine/100 g BW; maintenance doses: 5 mg ketamine/100 g BW, 0 mg xylazine) and placed on a stereotaxic platform inside an anechoic chamber.

The auditory stimulation consisted of 50-ms pure tone (100, 125, 150, 175, and 200 Hz sinusoidal frequency) pulses with 10-ms ramp up/down durations and 200-ms inter-stimulus intervals, delivered by a pair of multi-field magnetic speakers (MF1, Tucker-Davis Technologies, FL). Neural activities of MSO neurons were recorded using the sapphire optrode with LED illumination alternating at a 2-second interval. The raw recording traces from the optrode were pre-amplified using a custom-made 8-channel bio-amplifier board and sent to the data acquisition system (NIDAQ, National Instruments, TX) at a sampling rate of 50 kHz. After each recording, semi-automatic spike sorting based on Gaussian-mixture clustering was performed on band-pass filtered (300-5000 Hz, 4th order Butterworth) raw traces to extract single-unit activities.

## Acknowledgements

This research was funded by The University of Macau (MYRG2022-00111-IME), funded by The Science and Technology Development Fund, Macau SAR (SKL-AMSV(UM)-2023-2025), and supported by the joint funding of the Nature Science Foundation of China (NSFC) & the Macao Science and Technology Development Fund (FDCT) of China (Grant No. 62061160368 and 0022/2020/AFJ)). The authors would also like to acknowledge the financial support of the ZUMRI-Lingyange Semiconductor Joint Lab (CP-031-2022) and the Lingyange Semiconductor Incorporated, Zhuhai (CP-017-2022) and the Blue Ocean Smart System (Nanjing) Limited (CP-003-2023). The optogenetic in-vivo recordings were supported by NIH / NIDCD grant R01 DC 18401 to A.K.

## Author contributions

Yanyan Y.X. and B.-Z. L. contributed equally to this work. Y.X. was responsible for the hardware design, experimental design, performance characterization, data analysis, and manuscript writing. B.-Z. L. was responsible for designing and conducting the experiments, analyzing and visualizing the data, and contributing to the manuscript writing. X.H., Y.L., Z.L., X.Y., and L.L. performed the experiments. M.R. carried out the experiments and assisted in manuscript writing. L.W. and Yu Y.X. designed the hardware and were involved in data collection. Y.H. performed additional experiments. A.K., S.H.P., Z.Y., and T.C.L. contributed to the study’s conceptual development, hardware design, experimental design, and manuscript preparation. B.Z. supervised and directed the overall research effort.

## Financial disclosures/conflicts of interest

The authors declare no financial conflicts of interest or disclosures relevant to this work.

---

**Supplementary Figure 1.**
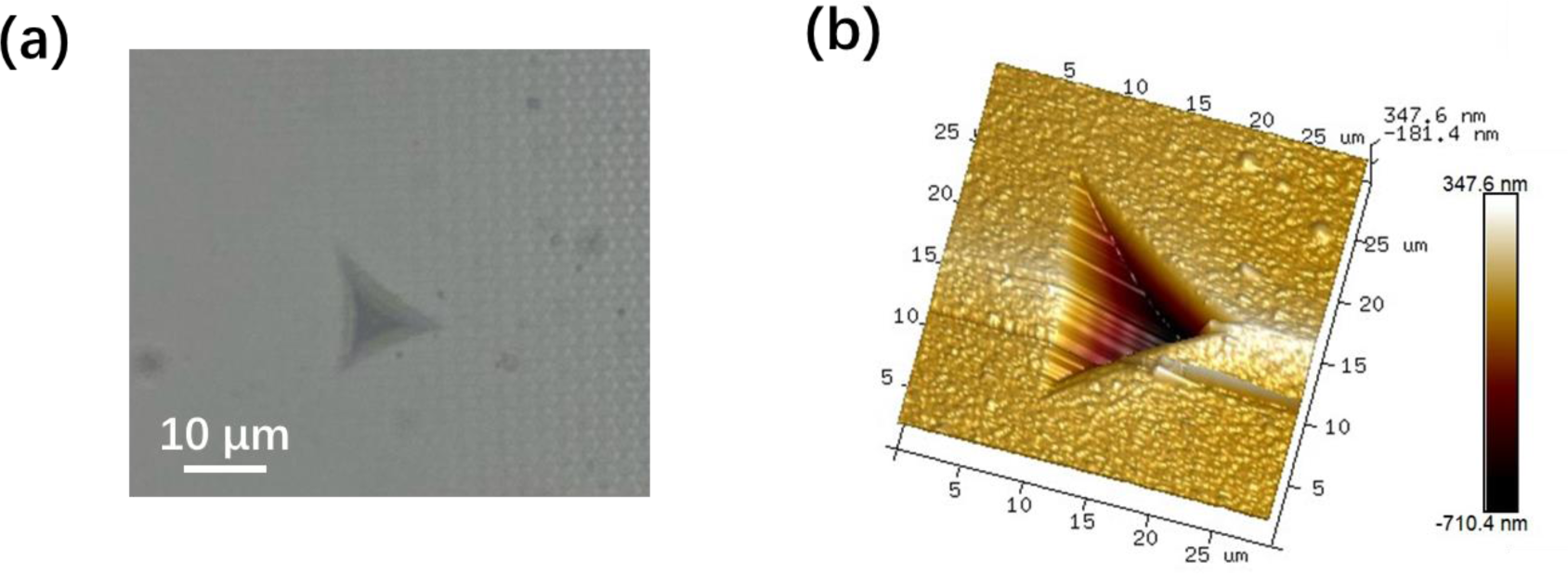
(a) Optical microscope image of the indentation morphology; (b) 3D AFM scanning image of the residual indentation.

## Notes

### Competing Interest Statement

The authors have declared no competing interest.

